# Dinophyceae use exudates as weapons against the parasite *Amoebophrya* sp. (Syndiniales)

**DOI:** 10.1101/2021.01.05.425281

**Authors:** Long Marc, Marie Dominique, Szymczak Jeremy, Toullec Jordan, Bigeard Estelle, Sourisseau Marc, Le Gac Mickael, Guillou Laure, Jauzein Cécile

**Affiliations:** IFREMER, Centre de Brest, DYNECO Pelagos, F-29280 Plouzané, France; UMR 7144 Sorbonne Université & Centre National pour la Recherche Scientifique, «Adaptation and Diversity in Marine Environment», Team «Ecology of Marine Plankton, ECOMAP», Station Biologique de Roscoff, 29680 Roscoff, France

## Abstract

Parasites of the genus *Amoebophrya* sp. are important contributors to marine ecosystems and can be determining factors in the demise of blooms of Dinophyceae, including microalgae commonly responsible for toxic red tides. Yet they rarely lead to the total collapse of Dinophyceae blooms. The addition of resistant Dinophyceae (*Alexandrium minutum* or *Scrippsiella donghaienis*) or their exudate into a well-established host-parasite culture (*Scrippsiella acuminata*-*Amoebophrya* sp.) mitigated the success of the parasite and increased the survival of the sensitive host. Effect were mediated via water-borne molecules without the need of a physical contact. Severity of the anti-parasitic defenses fluctuated depending on the species, the strain and its concentration, but never totally prevented the parasite transmission. The survival time of *Amoebophrya* sp. free-living stages (dinospores) decreased in presence of *A. minutum* but not of *S. donghaienis*. The progeny drastically decreased with both species. Integrity of the membrane of dinospores was altered by *A. minutum* which provided a first indication on the mode of action of these anti-parasitic molecules. These results demonstrate that extracellular defenses are an effective strategy against parasites that does not only protect the resistant cells but also have the potential to affect the whole surrounding community.

## Introduction

Parasites, which are supposed to account for half of the species richness, could make up the unseen majority of species extinctions (Carlson *et al*., 2017). Majority of parasites have essential ecological roles by contributing to the resilience of ecosystems, limiting the invasion and emerging of infectious diseases, and being critical to the biomass transfer between trophic levels (Johnson *et al*., 2013; Dougherty *et al*., 2016; Paseka *et al*., 2020). In marine ecosystems, parasites have a predominant role in the planktonic protists interactome inferred by sequence-based correlation networks (Lima-Mendez *et al*., 2015) and can represent up to 18% of interactions (Bjorbækmo *et al*., 2020). Parasites are important contributors to phytoplankton mortality and can even sometimes induce the demise of microalgal blooms (Brussaard, 2004; Chambouvet *et al*., 2008; Vardi *et al*., 2009). Amongst marine parasites, the Syndiniales *Amoebophryidae* (also called marine Alveolate group II, or MALVII) is a widely distributed family (Guillou *et al*., 2008; de Vargas *et al*., 2015), ubiquitous in marine waters, including ultra-oligotrophic ones (Siano *et al*., 2011), and has been associated with the demise of toxic species (Park *et al*., 2002; Chambouvet *et al*., 2008; Velo-Suárez *et al*., 2013; Li *et al*., 2014; Choi *et al*., 2017) in enriched coastal environments. Its life cycle is characterized by a free-swimming stage (zoospores, also named dinospores) followed by two successive intracellular stages (trophont then sporont) that eventually kills the host and release hundreds of dinospores. Dinospores are highly specialized and short-lived flagellated unicellular forms that only survive in culture a few hours to few a few days (Park *et al*., 2002).

*Amoebophrya* spp. are specialist parasites that require a compatible host to complete their life cycle. The overall consistency in the host spectrum observed within different strains of the same species suggest a genetic determinism underlying their host specialization (Cai *et al*., 2020). Many factors can influence the parasitic population dynamic such as physical (e.g. temperature, water column depth, physical mixing) and chemical (e.g. nutrients) parameters (Anderson & Harvey, 2020). Optimal abiotic conditions for parasitic infection do not always induce the collapse of targeted dinoflagellate blooms, implicating complex biotic interactions as fundamental player in the success of the parasite (Anderson & Harvey, 2020). Modelling approaches also indicate that the parasitic control of dinoflagellate blooms strongly depends on the structure (e.g. cell densities, grazing of free-living stages of parasite stages, competition between cells) of the plankton community (Alves-de-Souza *et al*., 2015). The co-existence between resistant and sensitive hosts could affect the parasitic propagation through different mechanisms, including a dilution effect (Alves-de-Souza *et al*., 2015; Alacid *et al*., 2016) or cell signaling between species.

Mechanisms of the host resistance against parasites are poorly known. Different strategies have been described so far, including the production of resting stages (Chambouvet *et al*., 2011a; Pelusi *et al*., 2020), the production of anti-parasitic metabolites produced internally (Pouneva, 2006; Bai *et al*., 2007; Place *et al*., 2009; Rohrlack *et al*., 2013; Scholz *et al*., 2017), and sometimes released into exudates (Scholz *et al*., 2017). The release of anti-parasitic compounds (APC) is a strategy that can be classified within the more general term of allelopathy. Allelochemicals refers to any secondary metabolite exuded by a microalga that affect the growth of another co-occurring protist (Granéli & Hansen, 2006). Whether and how the release of APC can influence the dynamic of parasite is a still an opened question.

This study investigated whether co-occurring Dinophyceae, resistant to *Amoebophrya* sp., can affect the dynamics of a parasite infecting a sensitive Dinophyceae host. For that, the infection dynamics and the dinospore survival was monitored in the well-established parasitic couple *S. acuminata* (ST147) infected by *Amoebophrya* sp. (A25) (Farhat *et al*., 2020) in presence and absence of two resistant dinoflagellate hosts: *Scrippsiella donghaienis* and *Alexandrium minutum*. These dinoflagellate species were selected for several reasons: (1) they can form recurrent dense blooms (Chapelle *et al*., 2014, 2015; Klouch *et al*., 2016) and are potential competitors of *S. acuminata*, (2) they co-occur with *S. acuminata* and its parasites *Amoebophrya* sp. in the same estuaries (Guillou *et al*., 2008; Cai *et al*., 2020), (3) they are resistant to *Amoebophrya* sp. A25 (Cai *et al*., 2020) and (4) the species *A. minutum* is a producer of allelochemicals that can negatively affect competing protists (Long *et al*., 2018). The production of such compounds by the species *S. donghaiensis* has not been reported so far.

## Materials and methods

### Biological material

#### Origin of strains and culture conditions

All hosts and parasitic strains originated from coastal marine waters of the NE Atlantic Ocean (Supporting Information Table S1). The parasite *Amoebophrya* sp. strain A25 was routinely maintained using the sensitive *S. acuminata clade* STR1 (ST147) as host. Resistant dinoflagellates used in this study were *A. minutum* (strains CCMI1002, Am176, Da1257) and *S. donghaienis* (strain Sc39 sampled during an *A. minutum* bloom). Infected and uninfected host cultures were maintained in a medium prepared with seawater from the Penzé estuary (27 PSU of salinity), stored in the dark for several months before used, filtered on 0.22 µm, autoclaved and enriched by a modified F/2 medium (Guillard’s Marine Water Enrichment Solution, Sigma) and 5% (v/v) soil extract (Starr & Zeikus, 1993). Cultures used for the Experiment 3 were prepared using a different medium (K, Keller et al. 1987, seawater from Argenton at 35 of salinity) after acclimation of strains. In both cases, a final filtration using a 0.22 µm pore size filter was processed after addition of nutritive solutions, under sterile conditions. Stock cultures and experiments were performed under continuous light (90-140 µEinstein m^-2^ s^-1^, light bulb Sylvania Aquastar F18W/174 or EASY LED universal light 438 mm) at 21 ± 1-2°C. All experiments were performed with plastic flasks (CytoOne vented flasks in polystyrene).

Uninfected hosts were kept in exponential growth phase by diluting 5 volumes of stock culture into 8 volumes of fresh medium every 3-4 days. Infections were propagated by diluting 1:5 (vol:vol) of the infected culture into healthy hosts *S. acuminata* (ST147) every 3-4 days. Physiological state of uninfected microalgal cultures was routinely screened using a handeld Pulse Amplitude Modulation (PAM) fluorometer (Aquapen-C AP-C 100, Photon Systems Instruments, Drassov, Czech Republic). As a prerequisite, only healthy cultures with a maximum (dark-adapted) photosystem II quantum yield (F_V_/F_M_) above 0.6 were used in experiments.

#### Synchronization and collect of *Amoebophrya* dinospores

Density and infectivity of dinospores decrease rapidly (days) after their release (Supporting Information Table S2), therefore the use of freshly released dinospores helps to maximize infections in the flask. To produce freshly released dinospores, cultures of parasites were synchronized (unless specified) following the protocol detailed in protocole.io dx.doi.org/10.17504/protocols.io.vrye57w. During synchronization, infections were initiated with 3-days-old cultures of *Amoebophrya* from which dinospores were collected after a gentle separation from the remaining host cells (*S. acuminata* ST147) using gravity filtration through nylon filter (5 μm, Whatman). These dinospores were incubated with the exponentially growing host *S. acuminata* (strain ST147) using a 1:2 parasite:host (vol:vol) ratio. After 24 hours of incubation, infected hosts were collected by filtration through a 5-µm-nylon-filter, then resuspended in an equal volume of fresh media, in order to remove remaining free-living dinospores. Three days later, freshly liberated dinospores of the same age (i.e. synchronized) were separated from remaining hosts by filtration as described before. In prior experiments, no effect of dilutions on the dinospore survival over 24 hours was observed whether using fresh culture medium, exudates from the healthy host ST147, or exudates from ST147xA25 infected culture (Supporting Information Table S2). Hereafter, ST147 filtrate was used to adjust densities by dilution.

#### Preparation of microalgal filtrates

Exudates from exponentially growing microalgal strains were collected by filtration (0.2 µm, acetate cellulose membrane, Minisart) using a gentle pressure process using a syringe. Exudates were diluted in filtrate obtained from a growing culture of ST147 in order to adjust concentrations. In the present study, dilution of exudates is expressed as equivalent to the microalgal density (corresponding to the theoretical concentration of cells that would have been reached by the initial culture after a similar dilution). Diluted exudates were then immediately used for experiments.

### Cell counting methods

#### Flow-cytometry (FCM): cell count and membrane permeability

Cell densities and parameters (e.g. forward scatter, size scatter, fluorescence signals) were estimated using a flow cytometer equipped with 488 nm and 405 nm lasers. A FACSAria flow cytometer (Becton Dickinson) was used in experiments 1 and 2; a Novocyte Advanteon (ACEA Biosciences) was used in Experiment 3. Dinophyceae were detected according to their red chlorophyll autofluorescence using the 488 nm laser. Free-living (dinospores) and late stages of infection of *Amoebophrya* spp. emit a bright green autofluorescence when excited under blue-violet light (Kim *et al*., 2004; Kim, 2006; Chambouvet *et al*., 2011a), a proxy of the parasite survival (Coats & Park, 2002). This natural autofluorescence was used to estimate the density of viable dinospores by FCM using the 405 nm laser. Intact cell membranes are impermeable to the SytoxGreen (SYTOX Green nucleic acid stain, Invitrogen), which only penetrates in cells having altered (i.e. permeable) membranes (DNA is then stained, and this autofluorescence is detected by FCM using the 488 nm laser). The SytoxGreen (final concentration of 0.05 μM) was incubated during 20 min in the dark before processing the samples.

#### Prevalence of infections (CARD-FISH)

Samples for CARD-FISH were fixed with paraformaldehyde (1% final concentration) for 1 hour at 4°C in the dark before filtration on a 0.8 µm, polycarbonate filter with a vacuum pump (< 200 mm Hg). Filters were then dehydrated using successive 50%, 80% and 100% ethanol solutions, dried and stored in the dark at −20°C. FISH-staining was then performed according to (Chambouvet *et al*., 2008). The prevalence was estimated under microscopy with an Olympus BX-51 epifluorescence microscope (Olympus Optical) equipped with a mercury light source, a 11012v2-Wide Blue filters set (Chroma Technology, VT, USA) and, with fluorescence filter sets for PI (excitation: 536 nm; emission: 617 nm) and FITC (excitation: 495 nm; emission: 520 nm).

Prevalence was determined by averaging infection counts on a minimum of 80 cells per replicate. Prevalence was divided in 3 categories: non-infected host cells, early stage (one or more dinospores of *Amoebophrya* sp. are found in the cytoplasm) and advanced stages (intermediate and beehive stages) as described in (Kayal et al., 2020). The progeny (i.e., the number of dinospores per infected host) was estimated by dividing the maximum concentration of dinospores by the concentration of infected host at advanced stages.

### Experimental set-ups

#### Experiment 1: Co-cultures

The dynamic of infection in co-cultures was compared when mixing the sensitive host (*S. acuminata* ST147) with the parasite (*Amoebophrya* sp. A25) and one resistant host (*A. minutum* CCMI1002 or *S. donghaienis* Sc39). Mixtures were prepared in triplicates, using a ratio *of 1:1:1*, with initial concentrations of 4000 cells mL^-1^ for each strain. Controls consisted of flasks containing (i) only the compatible host ST147 at 4000 cells mL^-1^ or (ii) the host (ST147) at 4000 cells mL^-1^ and its parasite A25 at a ratio of 1:1. An additional control consisted of mixing the host ST147 and one of the resistant host (CCMI1002 or Sc39) in parallel, replacing the parasite with 0.2 µm filtrate of the host culture. All cultures and controls were started simultaneously, using the same mother cultures. Cell densities were quantified once to twice per day by FCM. At the end of the experiment, samples were fixed with non-acidic Lugol’s solution (1% final concentration) for microscopic counts and differentiation between *S. accuminata* and *A. minutum* cells.

### Experiments 2 and 3: Evaluation of the effects of Dinophyceae filtrates on

#### Amoebophrya

Filtrates of microalgal cultures were used to analyze the effects of Dinophyceae exudates (from either *A. minutum* or *S. donghaienis*) on *Amoebophry*a. The Experiment 2 was organized into two parts, the first one to estimate the effect of Dinophycae exudates on the abundance of fluorescent dinospores over time, then the second to analyze their potential to successfully infect and produce a second generation of dinospores after 6 hours of contact with the filtrates.

First, dinospores from *Amoebophrya* sp. (A25) were exposed to serial dilutions of dinoflagellate filtrates (equivalent to 1000 and 5000 and 10000 cells mL^-1^) collected from three strains of *A. minutum* (Da1257, Am176, CCMI1002) and for one strain of *S. donghaienis* (SC39). Densities of fluorescent dinospores were monitored by FCM. The mortality rate (h^-1^) of autofluorescent dinospores was calculated over the first 3 hours according to equation 1, where N_1_ and N_2_ are the respective densities of autofluorescent dinospores before and after 3 hours of exposure to the filtrates. Controls consisted in the incubation of dinospores with exudates from the host ST147. Incubations for controls and using the highest filtrate concentrations (10000 cells mL^-1^) were performed in triplicates, while only one replicate was performed for intermediate exudate concentrations.

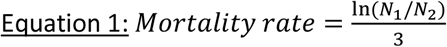

Then, dinospores previously exposed to the maximal concentration of exudates and in the control conditions after 6 hours of incubation were used for the second part of the experiment. Exposed-dinospores were mixed to the host strain ST147 at a dinospore:host ratio of 5:1 for dinospores exposed to *A. minutum* filtrates, and at three different ratios (0.5:1, 1:1, and 5:1) for dinospores exposed to *S. donghaienis* filtrate. The production of dinospores was monitored twice per day during 5 days by FCM, and the prevalence was analyzed after 47 hours of incubation by CARD-FISH in the controls and with the CMMI1002 and Sc39 filtrate treatments.

Experiment 3 was performed to monitor the concentrations of fluorescent dinospores and their membrane integrity over time when mixed with *A. minutum* exudates compared to the control. Dinospores from three days-old parasite cultures (non-synchronized) of *Amoebophrya* sp. A25 were harvested by filtration (5µm, cellulose acetate, Minisart). Dinospores were exposed in triplicate to *A. minutum* CCMI1002 filtrate at a final concentration of 5000 theoretical cells ml^-1^ in six well plates (CytoOne, made of polystyrene). In the control, dinospores were diluted in triplicate with *S. acuminata* (ST147) filtrate. The dinospore concentrations and the permeability of their membranes was estimated after 20, 40, 60 and 120 min of incubation with the filtrate.

### Statistics

All statistical analyses were performed using R software (R Foundation for Statistical Computing, Vienna, 2011). Significant differences in the different endpoints (e.g. concentrations of microalgae, concentrations of dinospores, prevalence) were assessed with a test of student or one-way ANOVA followed by a post-hoc Tukey HSD (ANOVA-HSD) when meeting the homoscedasticity with a Bartlett test and normality with a Shapiro-Wilk test. When homoscedasticity or normality could not be met, a non-parametric Krukal-Wallis test followed by a post-hoc Conover with a bonferroni adjustment was applied (KWbf). All tests were performed with a significance level of p-value = 0.05.

## Results

### Infections were mitigated by the presence of a resistant host

Experiment 1 tested whether the co-presence of a resistant host (*A. minutum* or *S. donghaienis*) could modify the *Amoebophrya* infection dynamics on a sensitive host (*S. acuminata*). In controls and when using fixed experimental culture conditions, a complete infection cycle lasted at least 51 hours, and ended with the sudden released of freshly produced dinospores (Fig. 1). During that period, infected host cells do not divide (Park *et al*., 2002), which explain the lower net growth rates recorded 25 hours after the parasite inoculation compared to the controls. Addition of a resistant host (CCMI1002 or Sc39) did not modify the duration of the parasite development, but always resulted in a significant decrease (> 60 %) of the dinospore production (Fig. 1). This observation could result from a deleterious effect on the sensitive host, a direct effect on the dinospore survival/infectivity, or both. Co-cultivation with *A. minutum* has a cost for *S. acuminata*. At the end of the experiment, densities of *S. acuminata* in the co-culture without parasite was of 6900 ± 1400 cells mL^-1^ while it reached 20000 ± 3000 cells mL^-1^ in the control.

**Figure 1:**
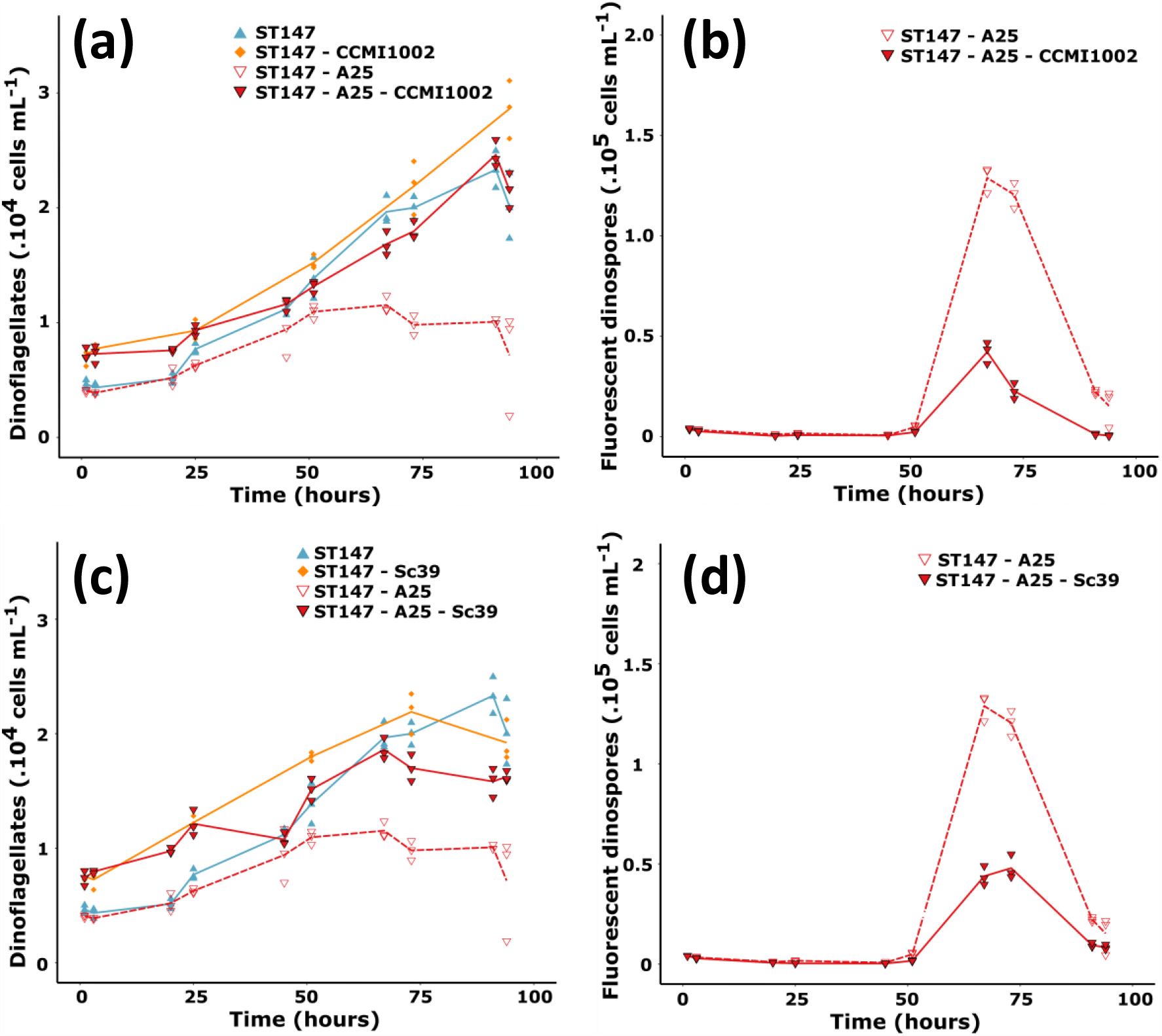
Co-cultures of *Amoebophrya* sp. (A25) with its compatible host *S. acuminata* (ST147) and a secondary resistant host, either *A. minutum* CCMI1002 (a, b) or *S. donghaienis* Sc39 (d, d). Densities of dinoflagellates (*S. accuminata* ST147 with *S. donghaienis* Sc39 or *A. minutum* CCMI1002) are shown in (a) and (c). Densities of fluorescent dinospores are shown in (b) and (d). Mean values are represented by lines, while replicate values are shown by the symbols. The same controls (ST147 and ST147-A25) are shown for both species as experiments were performed at the meantime.

### Exudates from *A. minutum* decreased the density of autofluorescent dinospores

Natural autofluorescence of dinospores can be used as a proxy for their viability (Coats & Park, 2002). In controls, 25% of autofluorescent dinospores were lost after 6 hours, leading to a natural mortality rate of 0.07 ± 0.01 h^-1^ in tested cultures conditions (Fig. 2). Experiment 2 tested whether the resistant dinoflagellate exudates affected this mortality rate. If no significant effect using *S. donghaienis* (Sc39) filtrate was observed, exposure to *A. minutum* filtrates resulted in a significant over-mortality (p-values < 0.02) compared to the control. This effect was however strain dependent, as illustrated by differing mortality rates between *A. minutum* strains (i.e. mortality of 0.11 ± 0.01, 0.92 ± 0.02 and 1.00 ± 0.01 h^-1^ of fluorescent dinospores after 6h of exposure using DA1257, AM176, and CCMI1002, respectively). After six hours, this exposure resulted in losses of 32 ± 1, 96.1 ± 0.2 and 97.2 ± 0.4 % of the initial density of fluorescent dinospores after 6h of exposure.

**Figure 2:**
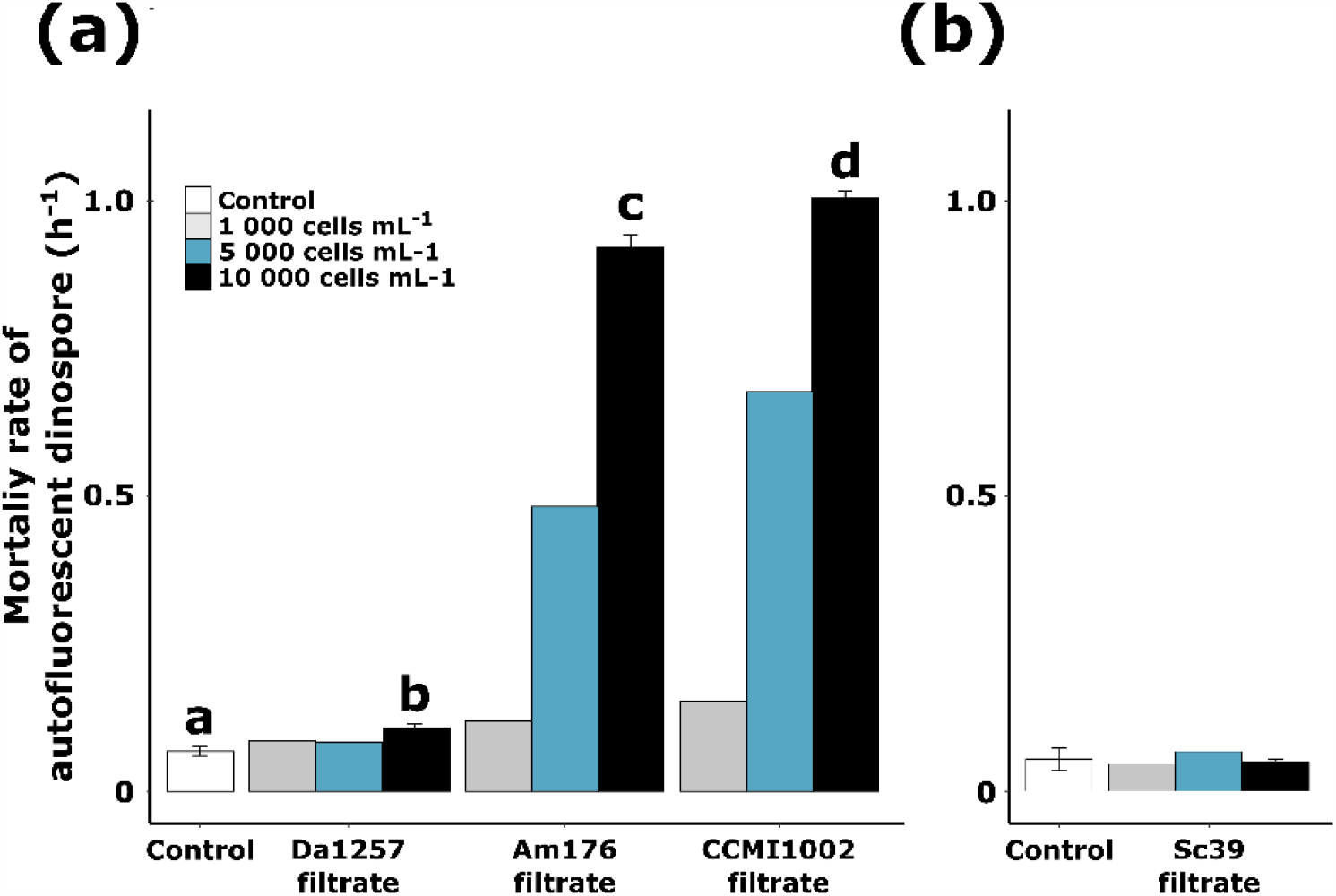
Maximal mortality rate of autofluorescent A25 dinospores in the different conditions. Dinospores were exposed to (a) *A. minutum* and (b) *S. donghaienis* filtrates during two separate sets of experiment. Results are expressed as the value or the mean ± standard deviation when triplicates were performed. Significant differences in the mortality rates are indicated by different letters. The complete dataset, with all sampling points (after 1, 3 and 6 hours) is provided in Supporting Information Fig. S1.

### Exudates from*A. minutum* decreased *Amoebophrya* sp. infectivity

To test whether the loss of fluorescence (Experiment 2) was linked to a loss of infectivity, dinospores were challenged for six hours with expsoure to exudates coming from three strains of *A. minutum* to fresh healthy host cultures. Densities were fixed for all treatments before the addition of exudates. However, because of the difference in mortality rates, the starting concentration of fluorescent dinospores differed over treatments (41000 ± 1400 dinospores mL^-1^ in the control, and 36000 ± 800, 2100 ± 100, and 1500 ± 200 dinospores mL^-1^ with exudates of Da1257, Am176, and CCMI1002, respectively). The ability of the remaining autofluorescent dinospores to infect their host even at low and unfavorable ratios was then explored. Novel infections were still observed in all treatments (Fig. 3). Filtrates of *A. minutum* did not seem to affect the intracellular stage as new progeny was released after 48 hours and the duration of infection was similar over treatments. Progeny (dinospore production per infected host) was 100 times lower with CCMI1002 than in the control (Table 1). As a result of lower prevalence and lower progeny, the maximal dinospore concentration were drastically decreased in the CCMI1002 and AM176 treatment as compared to the control or DA1257 filtrate (p-values < 10^−7^).

**Table 1 :**
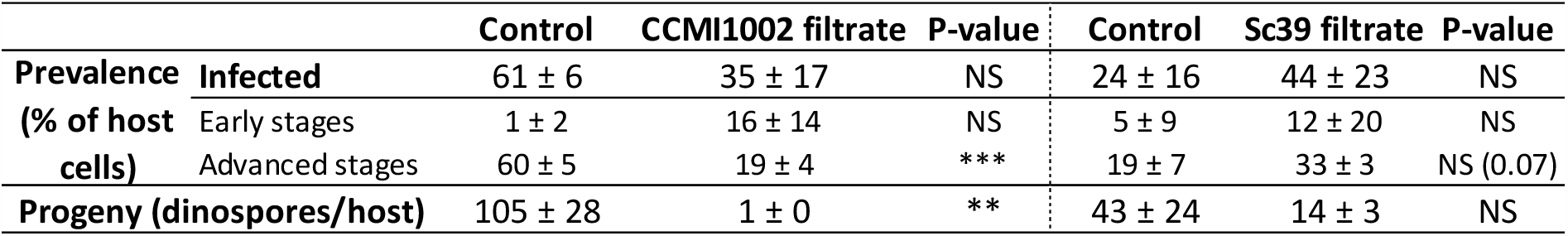
Prevalence of *Amoebophrya* sp. (A25) in *S. acuminata* (ST147) during Experiment 2 after 47 hours of contact. Two controls are shown as the two experiments were performed during two different sets. Significant values between the control and the dinophyceae treatment (CCMI1002 or Sc39) are indicated as followed: “NS” non significant, “*” 0.05> p-value > 0.01, “**” 0.01 > p-value > 0.001, “***” p-value < 0.001.

**Figure 3:**
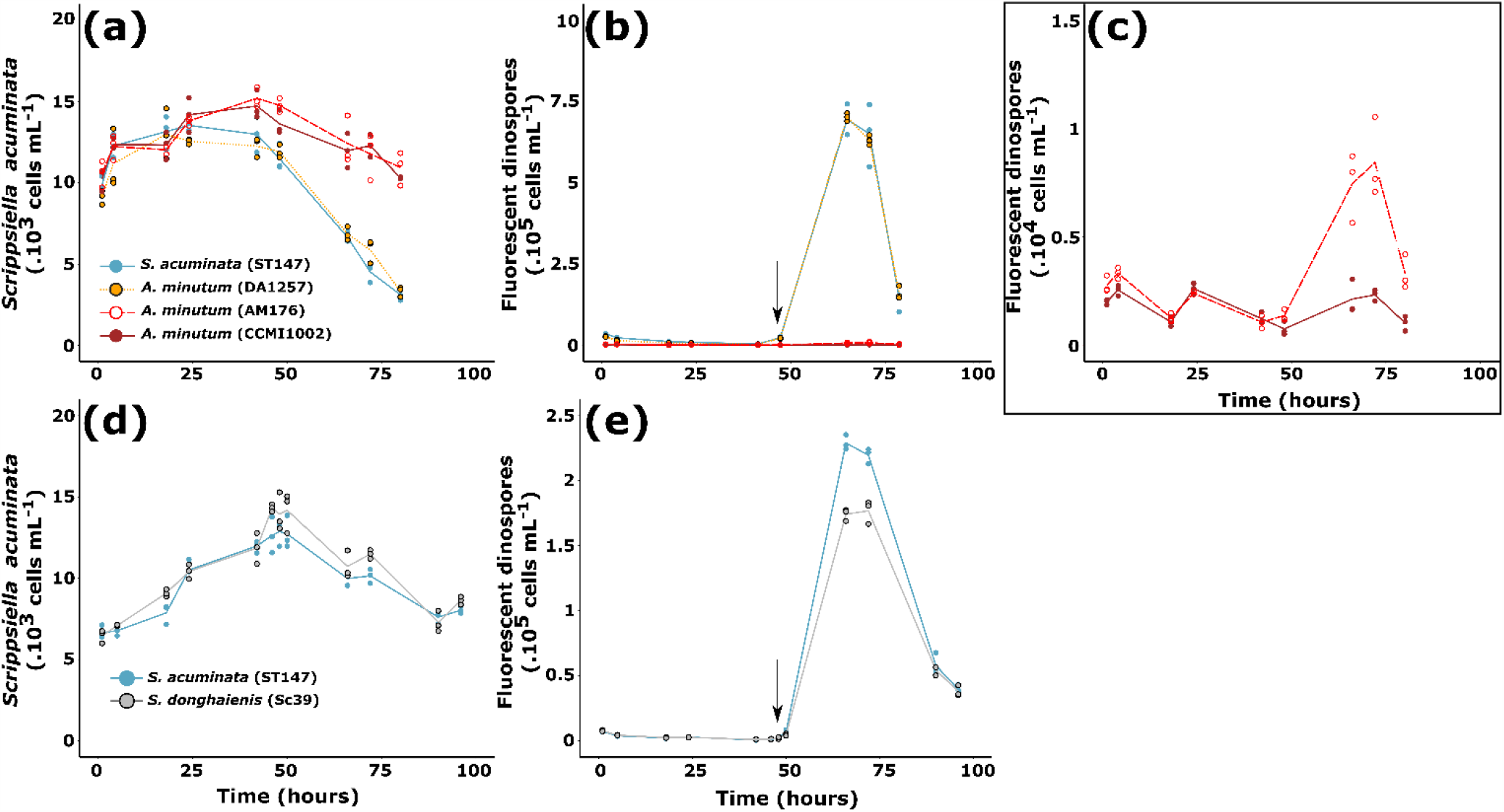
Effect of *A. minutum* (a-c) and *S. donghaienis* (d-e) filtrates (Theoretical cell concentration = 10^4^ cells mL^**-1**^) on infectivity of *Amoebophrya* sp. dinospores on its sensitive host *S. acuminata* (ST147). Cell densities of *S. acuminata* when mixed with A25 dinospores ar shown in (a, d). Dynamics of dinospores, previously exposed to the different filtrates, when mixed with the compatible host *S. acuminata* ST147 are shown in (b, c and e). (c) is a zoom of (b) with dinospores densities for Am176 and CCMI1002. *S. acuminata* (ST147; blue), *A. minutum* (Da1257; yellow), *A. minutum* (Am176; red) and *A. minutum* (CCMI1002; dark red). In experiments with *S. donghaienis* (Sc39; grey) filtrate, the graphs show results of the experiment at a dinospore: *S. acuminata* ratio of 1:1; results with other ratios can be found in Supporting Information Fig. S2. The arrow represents the sampling point for prevalence analysis. Lines represent the mean cell densities while the symbols represent the values of each replicate (N = 3).

The growth of the compatible host (*S. acuminata* ST147) was suppressed by the dinospores from the control or previously exposed to DA1257 filtrate. This suppression of the host growth in the control was linked to the high prevalence (61 ± 6% in the control; Table 1) of *Amoebophrya* sp. in host cells. In comparison, the compatible host in contact with the dinospores previously exposed to AM176 or CCMI1002 filtrates were still able to grow during the first 42 hours of incubation as the prevalence was lower (approximately a 35 % in the CCMI1002 treatment; Table 1). Between 42 and 80 hours, a collapse of the host population was observed in all conditions. The degree of the decline in host population was likely correlated to the prevalence of cells at advanced stages of infection (Table 1). With the CCMI1002 treatment, 30 ± 4% of host cell losses were estimated (Fig. 3) against 75 ± 2% of host cell losses in the control.

The same experiment was conducted with Sc39, results from ratio 1:1 is shown (Fig 3d,e), results from ratios 0.5:1 and 5:1 are available in Supporting Information Fig. S2. On contrary to *A. minutum* filtrates, infections started with the same density of fluorescent dinospores in the controls and in Sc39 treatments, as no effect was observed on the autofluorescence of dinospores. Filtrates of *S. donghaienis* did not seem to affect the intracellular stage as novel infections were observed and the duration of infection was similar to control conditions: the release of new progeny started between 48 and 50 hours (Fig. 3e). The previous treatment of dinospores with Sc39 filtrate did not significantly affected the prevalence of *Amoebophrya* sp. in the host population nor affected the growth rate of the host during the first 48 hours (Fig. 3d). With or without the previous treatment with *S. donghaienis* filtrate, a sharp decline of the host population, concomitant with release of new progeny, was observed after 48 hours. Overall there were no statistical differences in the percentage of lysed host cells between the treatments ST147 (37 ± 3%), and Sc39 (38 ± 4%). The main effect of pre-exposure of dinospores to Sc39 filtrate was observed on the new generation of dinospores: the treatment induced a significant decrease of 22% of the maximum concentration of the second generation of dinospores (Fig. 3e and Supporting Information Fig. S2). This decrease did not seem to be linked to a lower prevalence (Table 1) but was more likely related to a lower number of progeny per infected host, even though the 3 fold decrease was not statistically significant when compared to the control (Table 1).

### Exudates from *A. minutum* disrupted membranes of *Amoebophrya* sp

In Experiment 3, it was tested whether the loss of autofluorescence from dinospores is concomitant to the loss of membrane integrity when exposed to *A. minutum* filtrate. The most potent strain *A. minutum* (CCMI1002) was used during this experiment. Following the exposure, a rapid decrease in the density of auto-fluorescent dinospores was observed, with a 40% decrease within 20 min of exposure and a 98% decrease after two hours (Fig. 4). This loss of auto-fluorescent dinospores was preceded by the dinospore membrane permeabilization. After 20 min of exposure to the filtrate, 68% of the still auto-fluorescent dinospores were permeable to SytoxGreen.

**Figure 4:**
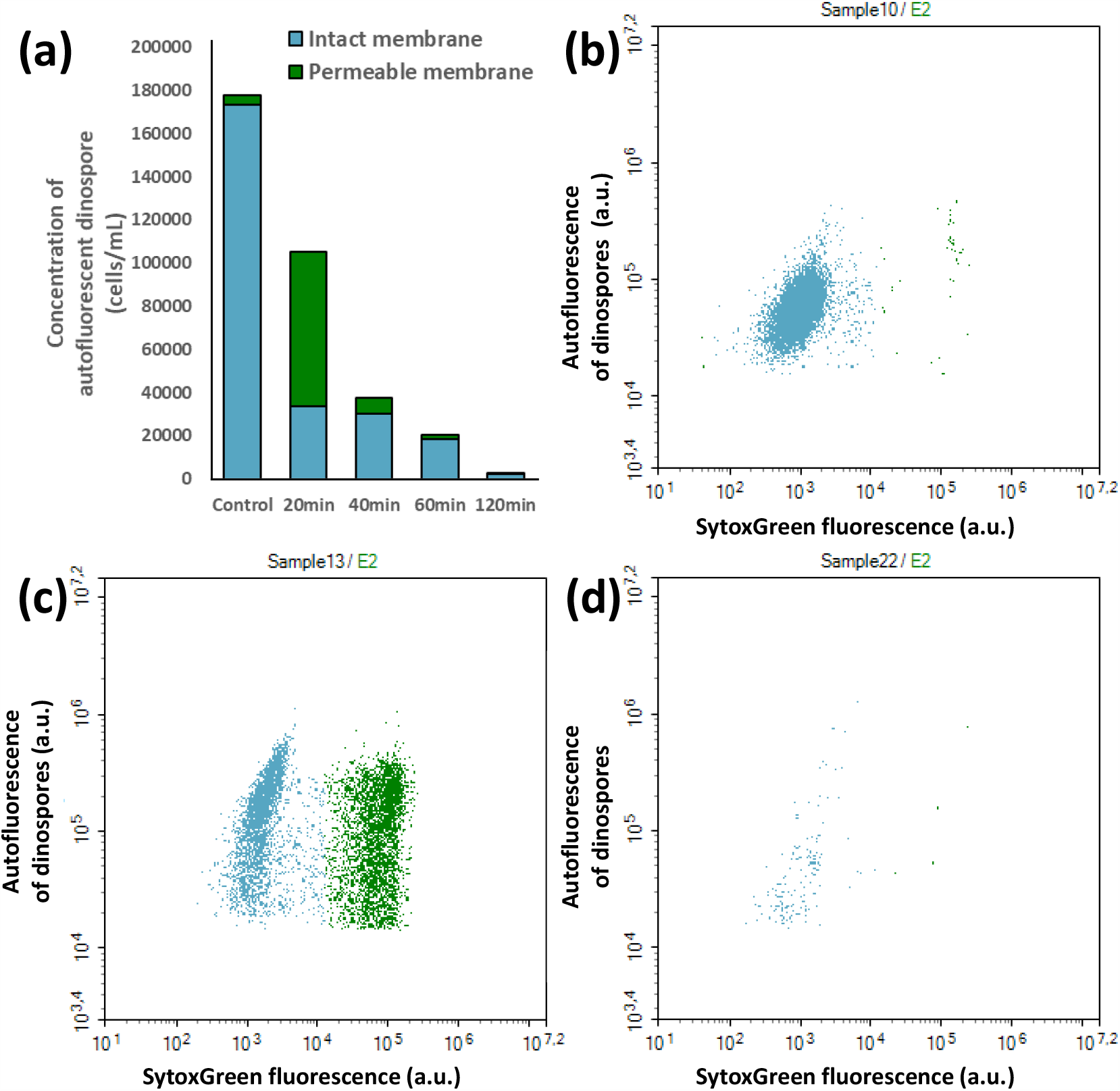
Effects of *A. minutum* filtrate on the density of auto-fluorescent (Green from 405 nm laser) dinospores and on the green fluorescence (from 488 nm laser) of cells after SytoxGreen staining. SytoxGreen stain only enters cells with damaged permeable membranes. (a) means of the cumulative densities (cells mL^**-1**^) of auto-fluorescent dinospores with impermeable (blue) and permeable (green) membranes to the stain. (b) dinospores in the control (stained but not exposed to *A. minutum* filtrate) and exposed to *A. minutum* filtrate for (c) 20 min, (d) 120 min.

## Discussion

Co-culture experiments with *A. minutum* showed that co-occurring resistant dinoflagellate species could either decrease survival of the free-living stage of the parasite, or limit its infectivity during the second generation, or both. Cells and filtrates of *A. minutum* caused similar effects to the infection dynamic, demonstrating that Dinophyceae can remotely affect parasites through the exudation of Anti-Parasitic Compounds (APC). Once released, APC are rapidly diluted, highlighting the importance of cell concentration and ratios. One may expect a particularly efficient protection for cells in close contact with the APC producers. Formation of dense cell patches with concentrations orders of magnitude higher than background (Durham & Stocker, 2012; Breier *et al*., 2018; Wheeler *et al*., 2019; Basterretxea *et al*., 2020) is likely more protective at micro-scales as this effect is density-dependent. As effects were observed using filtrates from cultures non-exposed to *Amoebophrya* sp. or its chemical cues, the release of APC is a passive defense mechanism. *A. minutum* exudates altered the integrity of the membrane prior the loss of the natural autofluorescence of the free-living stages of *Amoebophrya* sp. The loss of cell permeability might eventually lead to an osmotic cell lysis. The release of lytic APC by *A. minutum* cells in the phycosphere (i.e. microenvironment surrounding the cells (Seymour *et al*., 2017)) would act as a protective “shield” and must, at least partially, explain the resistance of *A. minutum* against *Amoebophrya* sp. This strategy was evidently ruled out by some *Amoebophrya* species, as it has already been reported that the genus *Alexandrium* could be infected by *Amoebophrya* sp. (Chambouvet *et al*., 2008; Lu *et al*., 2016). This could be explained by two hypotheses; (i) either *Amoebophrya* sp. only infects clones of *A. minutum* that do not release APC or, (ii) strategies to counteract APC effects exist. The second hypothesis has already been proven with another genus, *Amoebophrya* is able to acquire “antidotes” that enable it to avoid toxicity of *Karlodinium spp*. cells (Place *et al*., 2006), a potential host (Bai *et al*., 2007). This genus produces hydrophobic membrane permeabilizing compounds (Karlotoxins) with bioactivies and molecular targets that are similar to the permeabilizing compounds from *Alexandrium* (Ma *et al*., 2011; Long et *al*., Under review in *Harmful Algae*). The microalgal cells would be protected from their own toxins by their specific sterol membrane composition (Deeds & Place, 2006), a hypothesis also made for the genus *Alexandrium* and its allelochemicals (Ma *et al*., 2011). While cells from *Amoebophrya* sp. do not have a specific sterol signature (Leblond *et al*., 2006; Place *et al*., 2009), their sterol composition is rather related to sterol content of the host. The parasite is able to retain host lipid content, including the antidote for toxins, during the infection process. This strategy enables the parasite to avoid cell lysis and to infect its hostile host.

However, not all potential hosts are hostiles, the APC potency was highly variable between *A. minutum* strains. It was correlated with anti-microalgal (Long *et al*., 2018) and ichthyotoxic (Borcier *et al*., 2017; Castrec *et al*., 2018) activities. The mode of action of APC is very similar to the mode of action of anti-microalgal allelochemicals described from the same strain (Long *et al*., 2018; Long *et al*., Under review in *Harmful Algae*) and from *A. catenella* (formerly group I of the *A. tamarense/fundyense/catenella* species complex (Ma *et al*., 2011)). Both allelochemicals disrupt cell membranes and eventually induce cell lysis. It remains unclear whether APC are the same compounds than the ones described to have anti-microalgal or ichthyotoxic effects. However, it is also known that *Alexandrium* spp. can modulate its allelochemical potency against microalgae in response to changing physicochemical conditions (Martens *et al*., 2016; Long *et al*., 2019) but also its toxicity in response to cues from dead microalgal cells (Brown & Kubanek, 2020). It was also observed that genes associated with defensive responses such as the production of reactive oxygen species were overexpressed in a non-allelopathic strain of *A. fundyense* exposed to *Amoebophrya* sp. waterbone cues (Lu *et al*., 2016). Thus, activation and enhanced production of APC in the presence of a parasite cannot be excluded.

Similarly, *S. donghaienis* passively releases APC in the surrounding environment but a potential active defense remains to be investigated. In comparison with *A. minutum*, a different effect, potentially mediated by different molecules, was observed in the presence of *S. donghaienis*. The former species did not affect the survival of the free-living stage of the parasite infecting *S. acuminata*, but decreased its infectivity (i.e. ability to enter the cells) and/or progeny (i.e. ability to develop and produced the next generation of dinospores). The production of extracellular bioactive compounds was yet reported in *S. acuminata* (formerly identified as *S. trochoidea*; (Wang & Tang, 2008; Tang & Gobler, 2012) but never tested in *S. donghaeinis*. APC might also indirectly act as a signaling system for *S. acuminata* that could in turn modify its resistance against *Amoebophrya* sp., an interesting hypothesis that requires more investigation. Importantly, these results remind us that chemical weapons are not limited to harmful algal bloom species.

It was suggested that the presence of allelopathic genotypes could facilitate the proliferation of non-allelopathic cells and therefore the whole population (John *et al*., 2015; Felpeto *et al*., 2018). Here, it was additionally demonstrated that opportunistic (and competitive) species like *S. acuminata* could be protected from parasitism and could benefit from a few anti-parasitic producers among *A. minutum* and *S. donghaienis* populations. The cumulative protective effect provided by resistant hosts likely contributes to the survival of a sensitive dinoflagellate species in presence of its parasite, the private good becoming a public good (Driscoll *et al*., 2016). In cooperative associations, individuals that use a common goods produced by others in the absence of feedback are called cheaters. This is the case for non-allelopathic strains of *Prymnesium parvum* that benefits from the exclusion of diatom by another allelopathic strain (Driscoll *et al*., 2013). However, only the cheaters that are not or weakly sensitive to APC will benefit from the “cure”. For some microalgal species, the APC “cure” might have strong deleterious side effects. At least, a negative impact of *A. minutum* cells (but not of the filtrate) was observed on the growth of *S. acuminata* in co-cultures. After all, our results highlight a potential protective role of APC for the dinoflagellate but also suggest that the complexity of planktonic community structure in environmental communities may lead to contrasting results.

APC producers never induced the complete loss of the parasite, as illustrated by the production of a novel generation of dinospores, even in the presence of highly APC producers. These results suggest that once inside their host, the parasites might be totally or partially protected from APC. Eventually, such chemical defenses that moderate infections could contribute to the maintenance of the parasite in time, whilst avoiding the collapse of its hosts. More generally, allelopathy prevents competitive exclusion and promotes biodiversity in phytoplankton by favoring weaker competitors for nutrients (Felpeto *et al*., 2018). Similarly, APC promote biodiversity of parasites by favoring the most resistant parasite that may not be the most virulent. Indeed, these results well explain the discrepancies between the virulence of parasites that kill 100% of host cells within few days in the laboratory (this study and others;(Chambouvet *et al*., 2011a; Rodríguez & Figueroa, 2020)), and the coexistence of hosts and parasites in ecological studies that does not always result in the complete collapse of the sensitive population (Chambouvet *et al*., 2011b; Cosgrove, 2014). It also supports the hypothesis that the coexistence of different parasite cryptic species competing for the same host as reported by (Cai *et al*., 2020). All of these effects contribute to the explanation of the plankton paradox (Hutchinson, 1961). Chemical interactions between microorganisms tend to promote biodiversity (Czaran *et al*., 2002; Felpeto *et al*., 2018) and limit the effect of competitive exclusion for nutrients (or hosts for parasites) within plankton.

## Conclusion

Despite the ubiquity of genus *Amoebophrya* sp. in marine ecosystems, many opened questions remain on parameters affecting the parasite dynamic. This study highlight that resistant dinoflagellates can release exudates deleterious to the free-life stages of *Amoebophrya* sp. Chemical defenses must play a role in the resistance of dinoflagellates to parasites and more largely a role in their competitiveness. The exudation of anti-parasitic metabolites by resistant hosts in the surrounding environment provides a novel mechanistic link between a host-parasite couple and the surrounding community without the need of physical contact. The exudates not only protect the producer against parasitism but also have the potential to affect the whole community by decreasing the propagation of the parasite. This study revealed the importance of the composition of the plankton community during parasite infection as the severity of the effect fluctuated depending on the species and the strains of the resistant partner, their concentration and/or the ratio between the different partners. Another factor that has not been assessed in this study but requires further consideration is the potential of chemosensing in these interactions. Some parasites like the generalist parasite *Parvilucifera sinerae* can “sense” infochemicals (e.g. DMSP) from potential hosts (Garcés *et al*., 2013), even though they cannot actively select a compatible host (Alacid *et al*., 2016). Chemosensing of resistant host infochemicals by a parasite might have significant consequences on the efficiency of anti-parasitic defenses and should be studied through micro-scale studies. Overall, this study confirms that in the era of the -omic tools, “reductionist” experiments are still required to disentangle interactomes (Fitzpatrick *et al*., 2020). In addition to the ecological relevance, the use of anti-parasitic compounds extracted from dinoflagellates might be a mean to mitigate the parasites that can have devastating effects in algae mass cultures (Carney & Lane, 2014).

## Acknowledgements

This study was carried out with the financial support of IFREMER, the Centre National de la Recherche Scientifique (CNRS), Sorbonne Université, the Région Bretagne and the GDR Phycotox (a CNRS/IFREMER network on HABs). It results from different projects: the project PARALLAX (IFREMER), the project PARACIDE (SAD2018 Région Bretagne/IFREMER, GDR Phycotox) and the PRC France-Korea *MALV-REF*. The authors would like to warmly thank A/Prof. Dianne F. Jolley for the comments on the manuscript and English corrections.

## Author contributions

ML, LG, CJ, MS, MLG designed the study. ML, JS, DM, JT, CJ performed the experiments. ML, LG and CJ performed the data analysis and wrote the manuscript that was discussed and revised by all co-authors.

## Data availability

The data that supports the findings of this study are available in the supplementary material of this article.

## Supporting Information

**Table S2:**
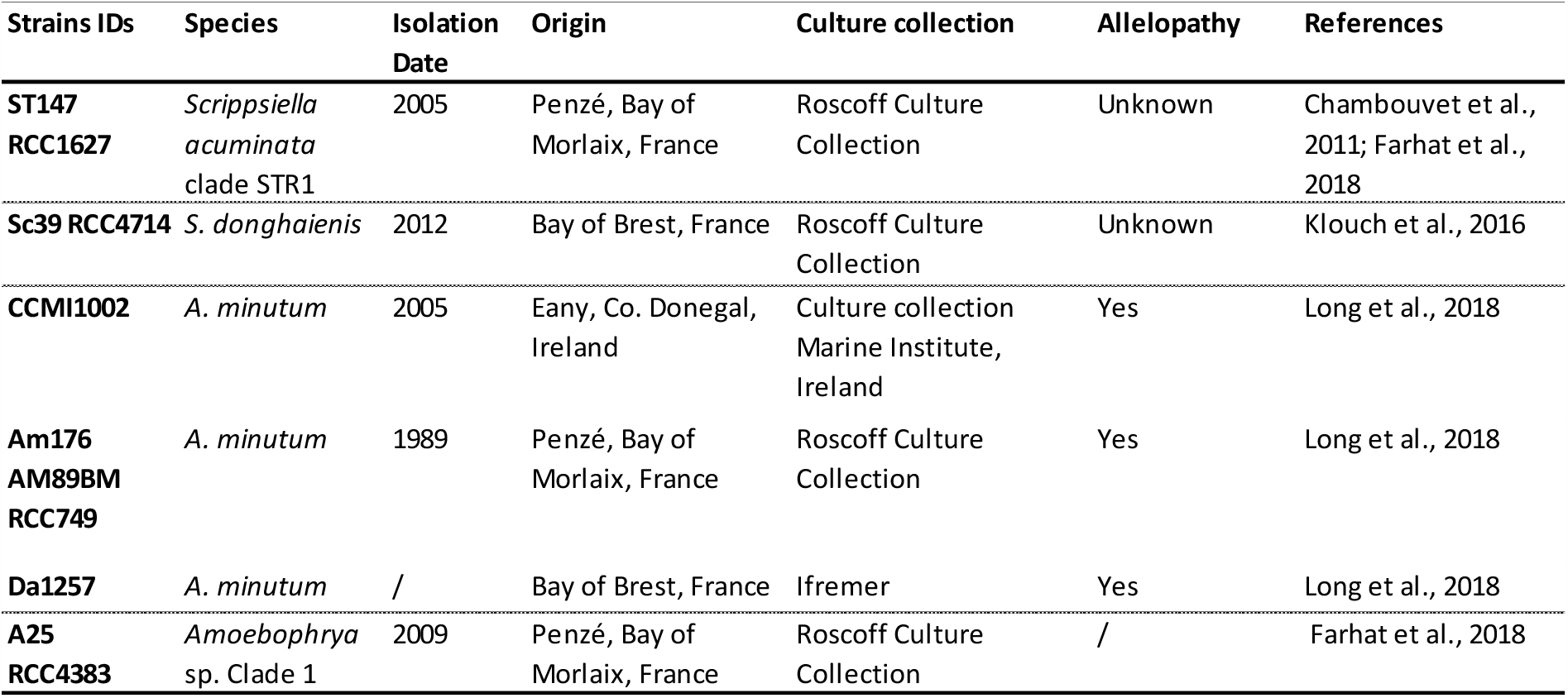
Details of microalgal and Syndiniales strains used in this study. “/” means that the data is unknown.

| Strains IDs | Species | Isolation Date | Origin | Culture collection | Allelopathy | References |
| --- | --- | --- | --- | --- | --- | --- |
| **ST147**<br>**RCC1627** | *Scrippsiella acuminata* clade STR1 | 2005 | Penzé, Bay of Morlaix, France | Roscoff Culture Collection | Unknown | Chambouvet et al., 2011; Farhat et al., 2018 |
| **Sc39 RCC4714** | *S. donghaiensis* | 2012 | Bay of Brest, France | Roscoff Culture Collection | Unknown | Klouch et al., 2016 |
| **CCMI1002** | *A. minutum* | 2005 | Eany, Co. Donegal, Ireland | Culture collection Marine Institute, Ireland | Yes | Long et al., 2018 |
| **Am176**<br>**AM89BM**<br>**RCC749** | *A. minutum* | 1989 | Penzé, Bay of Morlaix, France | Roscoff Culture Collection | Yes | Long et al., 2018 |
| **Da1257** | *A. minutum* | / | Bay of Brest, France | Ifremer | Yes | Long et al., 2018 |
| **A25**<br>**RCC4383** | *Amoebophrya* sp. Clade 1 | 2009 | Penzé, Bay of Morlaix, France | Roscoff Culture Collection | / | Farhat et al., 2018 |

**Table S2:** Effect of culture medium or filtrates on the mortality rate of autofluorescent dinospores of *Amoebophrya* sp. A25 over 24 hours. Loss rate was calculated according to Equation 1 in the manuscript. “NS” means that no significant difference was observed.

|  | Initial density | 24 hours density | Loss rate of autofluorescent dinospores | Significance |
| --- | --- | --- | --- | --- |
| **F/2 media** | 29500 ± 400 | 14000 ± 1000 | 0.031 ± 0.004 | NS |
| **ST147 filtrate** | 27000 ± 900 | 15000 ± 2000 | 0.026 ± 0.005 | NS |
| **A25-ST147 filtrate** | 29000 ± 1500 | 14000 ± 1000 | 0.030 ± 0.002 | NS |

**Figure S1 :**
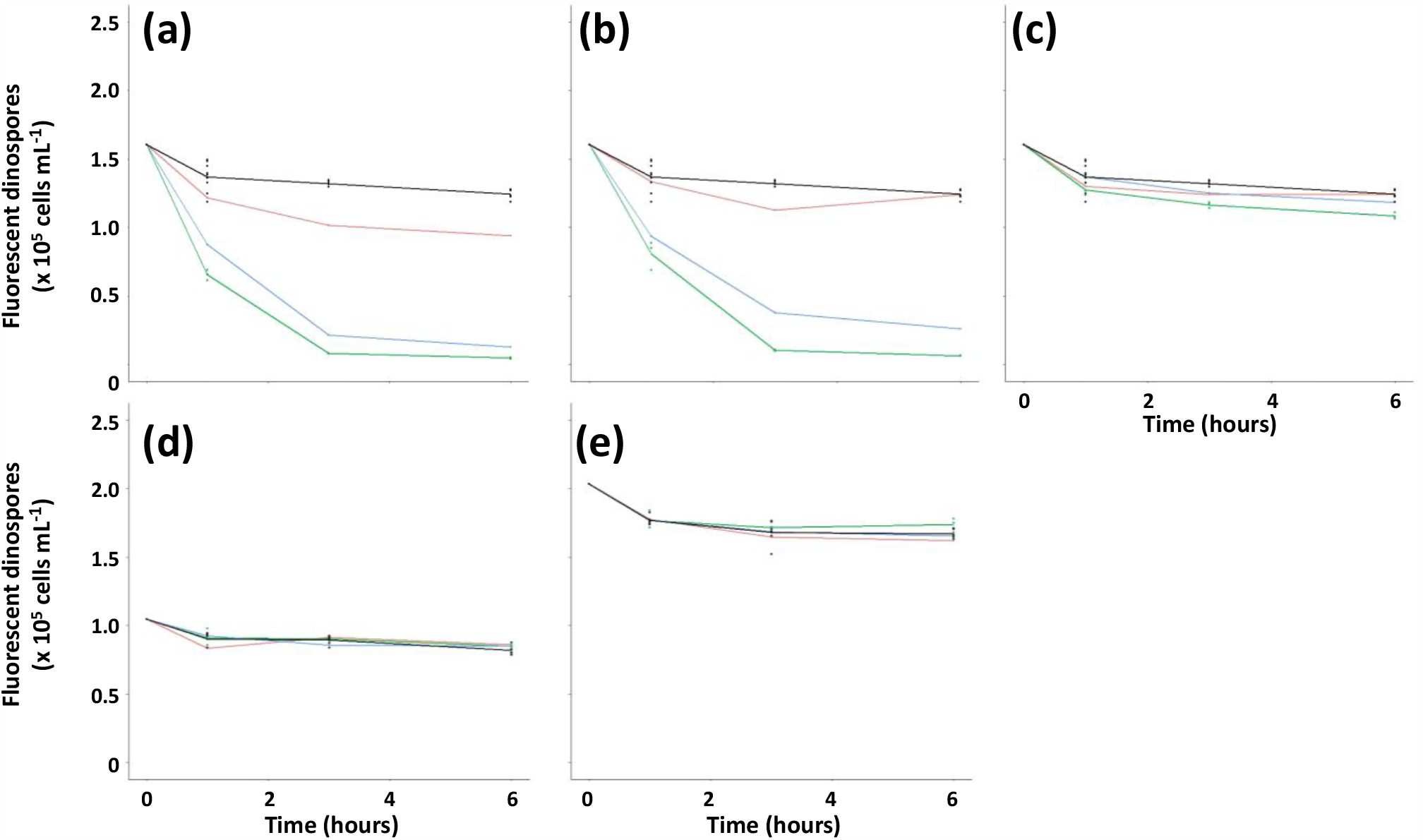
Concentration of fluorescent dinospores over 6 hours after exposure to (a) *A. minutum* CCMI1002, (b) *A. minutum* AM176, (c) *A. minutum* DA1257 and (d-e) *S. donghaienis* Sc39. Dinospores were exposed to ST147 filtrate (control in black), and filtrate at equivalent microalgal densities of 1000 (red lines), 5000 (blue lines) and the maximum concentration of (a, b, c, e) 10000 cells mL^-1^ and (d) 7000 cells mL^-1^ (green lines).

**Figure S2:**
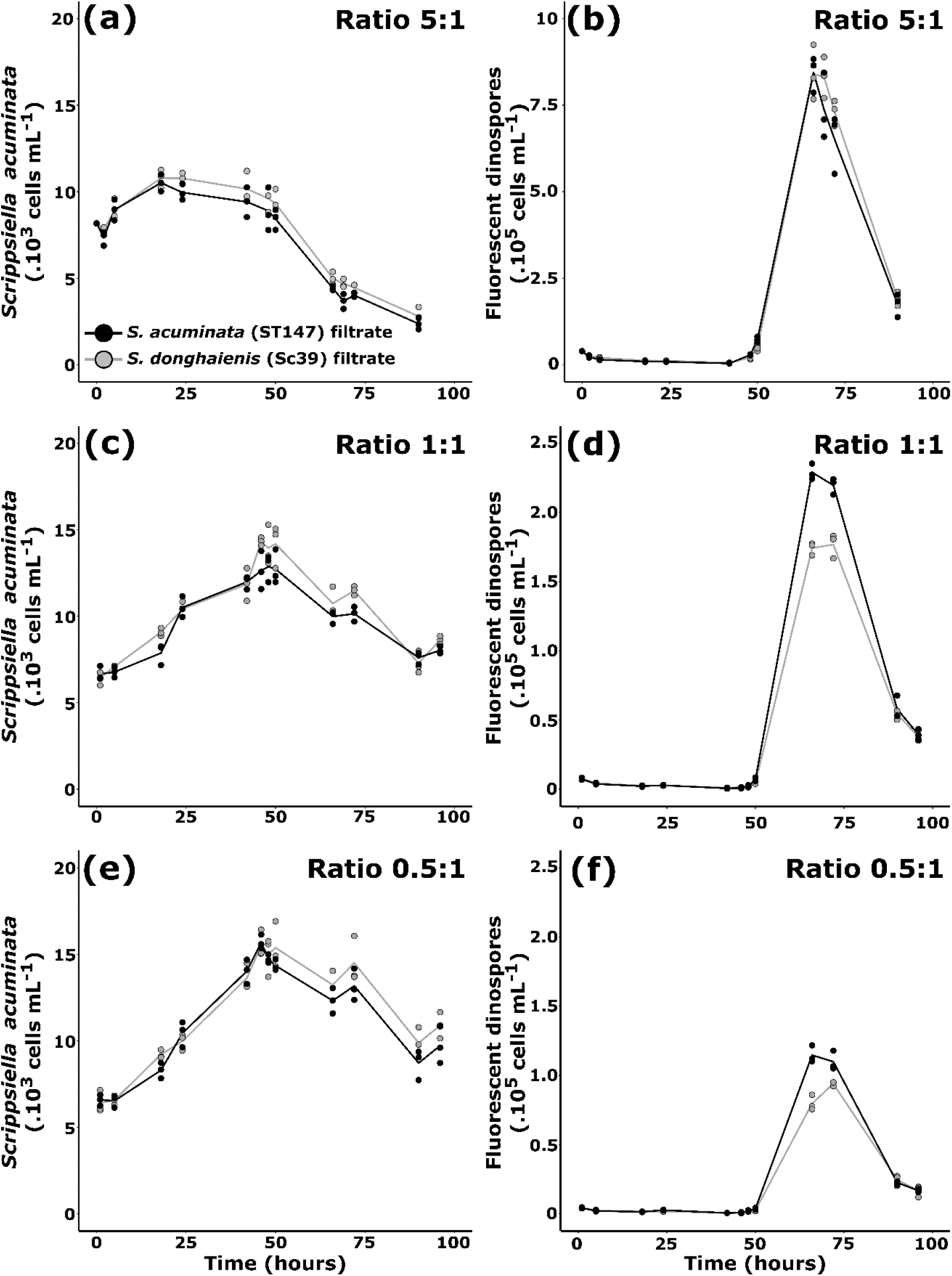
Effect of *Sc39 S. donghaienis* filtrates (Theoretical cell concentration = 7000 cells ml^**-1**^ for (a) and (b), 10000 cells mL^**-1**^ for (c, d, e, f) on the infectivity of A25 dinospores to *S. acuminata* (ST147). Infections were performed at three different dinospores: *S. acuminata* ratios; 5:1(a, b),1:1 (C and D) and 0.5:1 (e, f). The densities of the host ST147 during the infection cycle are shown in graphs (a, c, e). The density of fluorescent dinospores are shown in graphs (b, d, f). The controls (Filtrate ST147) are shown in black while the conditions in presence of Sc39 filtrate are shown in grey.

